# Within-family studies for Mendelian randomization: avoiding dynastic, assortative mating, and population stratification biases

**DOI:** 10.1101/602516

**Authors:** Ben Brumpton, Eleanor Sanderson, Fernando Pires Hartwig, Sean Harrison, Gunnhild Åberge Vie, Yoonsu Cho, Laura D Howe, Amanda Hughes, Dorret I Boomsma, Alexandra Havdahl, John Hopper, Michael Neale, Michel G Nivard, Nancy L Pedersen, Chandra A Reynolds, Elliot M Tucker-Drob, Andrew Grotzinger, Laurence Howe, Tim Morris, Shuai Li, MR within-family Consortium, Wei-Min Chen, Johan Håkon Bjørngaard, Kristian Hveem, Cristen Willer, David M Evans, Jaakko Kaprio, Bjørn Olav Åsvol, George Davey Smith, Bjørn Olav Åsvold, Gibran Hemani, Neil M Davies

**Affiliations:** K.G. Jebsen Center for Genetic Epidemiology, Department of Public Health and Nursing, NTNU, Norwegian University of Science and Technology, Norway; Medical Research Council Integrative Epidemiology Unit, University of Bristol, BS8 2BN, United Kingdom; Clinic of Thoracic and Occupational Medicine, St. Olavs Hospital, Trondheim University Hospital; Population Health Sciences, Bristol Medical School, University of Bristol, Barley House, Oakfield Grove, Bristol, BS8 2BN, United Kingdom; Postgraduate Program in Epidemiology, Federal University of Pelotas, Pelotas, Brazil; Netherlands Twin Register, Department of Biological Psychology, Vrije Universiteit Amsterdam, Amsterdam, The Netherlands; Nic Waals Institute, Lovisenberg Diaconal Hospital Spångbergveien 25, 0853 Oslo, Norway; Department of Mental Disorders, Norwegian Institute of Public Health, Sandakerveien 24C, 0473 Oslo, Norway; The University of Melbourne, 207 Bouverie St, Carlton, Victoria, 3010, Australia; Virginia Institute for Psychiatric and Behavior Genetics, Virginia Commonwealth University, Richmond, Virginia, USA; Karolinska Institutet, Dept of Medical Epidemiology and Biostatistics, Stockholm, Sweden; Department of Psychology, University of California Riverside, Riverside, CA, USA; Department of Psychology and Population Research Center, University of Texas at Austin, 108 E. Dean Keeton Stop A8000, Austin TX 78712 USA; Centre for Epidemiology and Biostatistics, Melbourne School of Population and Global Health, The University of Melbourne 207 Bouverie Street, Carlton, Victoria 3053, Australia; Centre for Cancer Genetic Epidemiology, Department of Public Health and Primary Care, University of Cambridge, Strangeways Research Laboratory, Worts Causeway, Cambridge CB1 8RN, United Kingdom; Center for public health genomics, Department of public health sciences, University of Virginia, Charlottesville, VA, USA; Department of Biostatistics and Center for Statistical Genetics, University of Michigan, Ann Arbor, USA; Department of Internal Medicine, University of Michigan, Ann Arbor, MI, US; Department of Human Genetics, University of Michigan, Ann Arbor, USA; University of Queensland Diamantina Institute, Translational Research Institute, Brisbane, Queensland, Australia; Department of Public Health, University of Helsinki, Helsinki, Finland; Institute for Molecular Medicine Finland (FIMM), University of Helsinki, Helsinki, Finland; Department of Endocrinology, St Olavs Hospital, Trondheim University Hospital, Trondheim, Norway

**Keywords:** Mendelian randomization, Gene-environment correlation, Population stratification, Assortative mating, Dynastic effects, Educational attainment

## Abstract

Mendelian randomization (MR) is a widely-used method for causal inference using genetic data. Mendelian randomization studies of unrelated individuals may be susceptible to bias from family structure, for example, through dynastic effects which occur when parental genotypes directly affect offspring phenotypes. Here we describe methods for within-family Mendelian randomization and through simulations show that family-based methods can overcome bias due to dynastic effects. We illustrate these issues empirically using data from 61,008 siblings from the UK Biobank and Nord-Trøndelag Health Study. Both within-family and population-based Mendelian randomization analyses reproduced established effects of lower BMI reducing risk of diabetes and high blood pressure. However, while MR estimates from population-based samples of unrelated individuals suggested that taller height and lower BMI increase educational attainment, these effects largely disappeared in within-family MR analyses. We found differences between population-based and within-family based estimates, indicating the importance of controlling for family effects and population structure in Mendelian randomization studies.

---

Mendelian randomization is an approach that uses genetic variants as instrumental variables to investigate the causal effects of one trait (the ‘exposure’) on another (the ‘outcome’).^1–5^ It has gained popularity due to the recent expansion in the scale of genome wide association studies (GWAS) and because it can ameliorate bias due to processes of residual confounding and reverse causation that affect most other observational approaches. In order for Mendelian randomization estimates to be valid, the genetic instrument must meet three assumptions: 1) relevance, it must associate with the exposure, 2) independence, there must be nothing that causes both the instrument and the outcome, and 3) exclusion, the association of the instrument and the outcome must be entirely mediated via the exposure. Methodologists have focused on developing methods to overcome bias in Mendelian randomization studies due to horizontal pleiotropy,^6–11^ which would violate the exclusion assumption. However, in this paper our attention is focused on the second assumption: independence. We demonstrate how population and family structures can lead to violations in the second assumption, and that traditional family-based methods are well placed to rectify this problem.

Mendel’s laws of genetic inheritance provide a rationale for why much genetic variation will be independent of the environment and genetic variation for other traits.^1,12^ However, environmental and social factors such as assortative mating, dynastic effects, and population structure may affect the distribution of genetic variants for specific traits within populations (see **Box 1** in supplement for definitions).^13–16^ **Figure 1** illustrates the impact of these processes in the context of Mendelian randomization, the commonality amongst all three processes being that they induce a spurious association between the instrumenting variant and the outcome through confounding. Assortative mating can occur when individuals select a partner based on a selected phenotype.^6,17^ For example, couples tend to have more similar age, education, and body mass index than would be expected by chance.^18,19^ If assortative mating arises due to individuals with a particular genetic predisposition selecting mates who have a particular genetically influenced phenotype, this can induce spurious genetic associations which can result in biased estimates from Mendelian randomization studies.^6^ In addition, social homogamy may lead to people selecting partners who are similar to themselves,^20^ which can compound across generations.^6^ Dynastic effects can occur when the expression of parental genotype in the parental phenotype directly affects the offspring phenotype. For example, higher educated parents might support their children’s education by providing a stimulating environment, being able to afford tutoring for their child, buying homes in better school districts, or paying for private schools. Finally, residual population structure occurs when there are geographic or regional differences in allele frequency relating to a trait of interest that cannot necessarily be controlled for via principal components.^13^ Confounding by population stratification,^1^ in which ancestry is correlated with both phenotypes and genotypes, was a major concern during early development of Mendelian randomization.^1^ However, this fear was gradually assuaged by a decade of GWAS results that were apparently reliable in the face of population structure.^21^ GWAS are now performed on a huge scale; as a consequence the problem of population stratification is again of potential concern because the high statistical power of large studies renders them susceptible to bias from very subtle population structure.^13,22^

**Figure 1.**
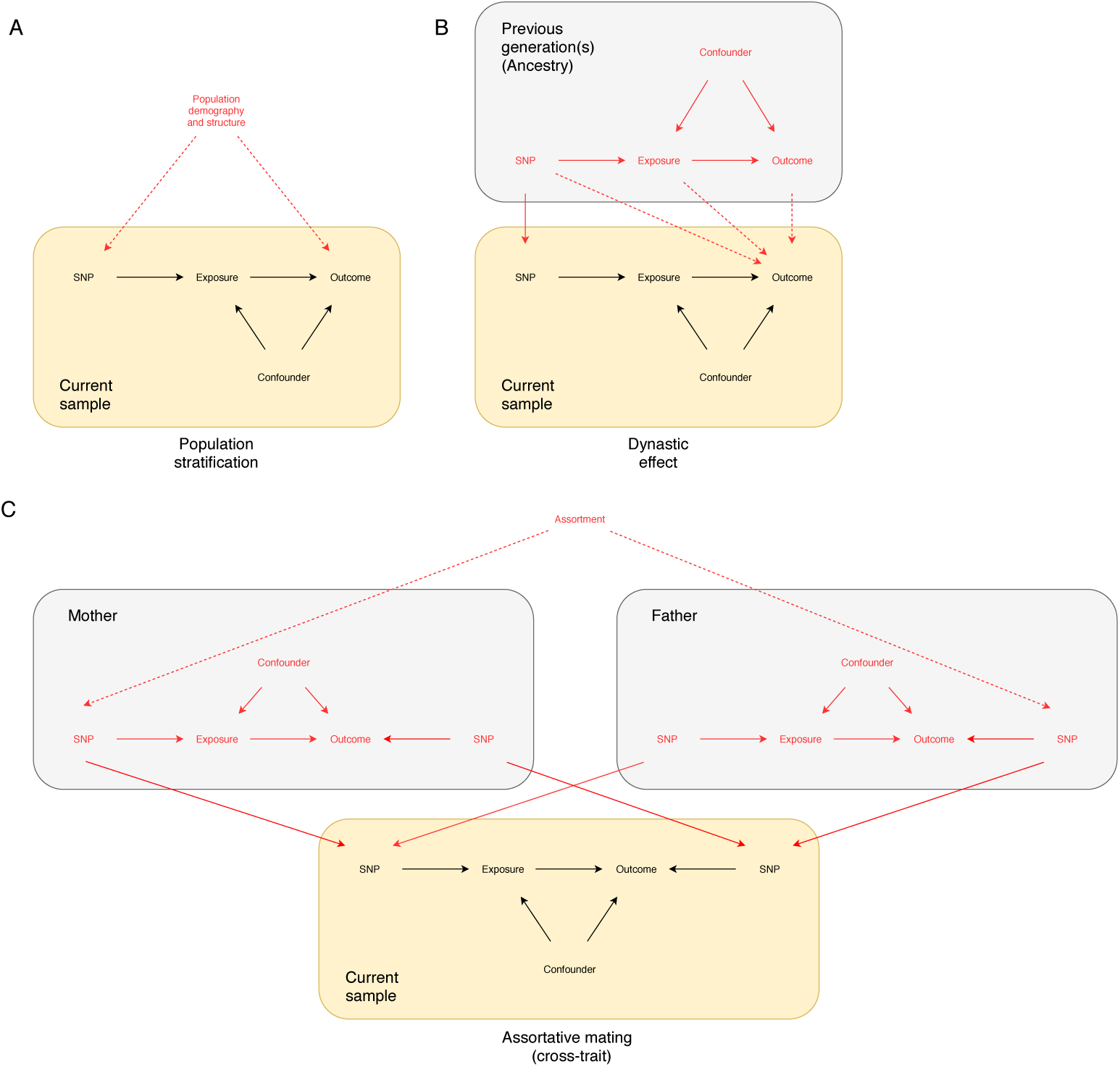
Dynastic effects can cause confounding bias in Mendelian randomization studies. The MR estimate of the causal effect of the exposure on the outcome is biased because of potentially unobserved confounders between the SNPs and the exposure and the outcome. Panel A illustrates how population demography and structure can confound the SNP-outcome association. Panel B illustrates how dynastic effects can confound the SNP-outcome association. The solid red vertical arrow indicates the genetic inheritance of germline DNA. The dotted line indicates the direct (dynastic) effect of the parents on the offspring’s outcomes. Mendelian randomization estimates of the effect of the exposure on the outcome in samples of unrelated individuals will be biased, because there is a path between offspring SNP and the outcome via the effect of the parents’ phenotypes on their offspring’s outcomes (dynastic effects). The presence of dynastic effects would violate one of three key Mendelian randomization (instrumental variable) assumptions – the independence assumption. Estimates that control for mother or father genotype, or sibling genotype will close this path and be unbiased. Panel C illustrates how assortative mating can confound the SNP-outcome association. In this example we present cross-trait assortative mating where there is a pathway between the mother’s genotype and offspring’s outcome via the father’s genotype for the outcome. Dynastic effects and assortative mating can be accounted for by using methods based on within-family contrasts.

### Box 1: Possible confounders of genetic associations

#### Assortative mating

when individuals choose their partners non-randomly, so are more alike or selected on a particular trait than would be expected. This can occur because people select on a specific trait, e.g. if tall women prefer to marry tall men, people who drink alcohol choose partners who also drink,^8^ or because of social homogamy where people select partners who have a similar environmental background to themselves, e.g. if educated women select men with a similar amount of education as themselves who happen to be taller.^71^

#### Dynastic effects

when parental genotype affect offspring outcomes through pathways other than via offspring genotype. For example, if more educated parents support their offspring’s education, or if parents smoking positively or negatively affected the likelihood of their offspring smoking. An example of dynastic effects are passive gene-environmental correlations.^11,16,17^

#### Fine scale population structure

when subtle difference in ancestry are associated with offspring phenotypes. For example, on average individuals from the north and west of England are poorer and are less educated, there are also geographic gradients in the distribution of education associated variants.^13^

Confounding in genetic association estimates, as induced by assortative mating, dynastic effects and population stratification, can and has been resolved by using family-based study designs.^6,23,24^ For example, in sibling pair studies, genetic associations at loci can be partitioned into between pair and within pair components.^23^ Because genetic differences within sibling pairs reflect random independent meiotic events, within pair effects are unrelated to population stratification and most potential confounders that might influence the phenotype. Similarly, other family-based designs and within-family tests to adjust for or exploit parental genotypes exist, such as estimating maternal and offspring genetic effects using structural equation modelling,^25^ quantitative transmission/disequilibrium tests,^26,27^ or mother-father-offspring trios to adjust for parental genotypes.^28^ Such within-family designs have been used to validate results from genome wide association studies,^29,30^ obtain unbiased heritability estimates,^31^ and assess causation in the classical twin design.^32,33^ Yet, despite the initial extended proposal of Mendelian randomization advising that the only way to ensure true randomization was through a within-family design^1^, to-date contemporary implementations using modern genomic methods have rarely been performed. The principal reason for this has been a lack of genomic data collected from families at a scale sufficient to be suitably powered. However, as we enter the age of national scale biobanks and very large twin studies, this essential extension of Mendelian randomization is becoming feasible.

This paper presents theory and simulations that demonstrate how within-family designs can be coupled with genomic data to perform unbiased Mendelian randomization analyses. We integrate these approaches in a modular fashion alongside other methods that have been developed for pleiotropy-robust inference (i.e. to be resilient to violations of the third assumption of Mendelian randomization).^7–9,34^ Using 28,777 siblings from HUNT and 32,231 siblings from the UK Biobank, we illustrate these methods empirically. First, we analyse the influences of BMI on high blood pressure and risk of diabetes as positive controls. Second, we evaluate the influences of height and BMI on educational attainment and demonstrate major discrepancies between population-based and within-family based approaches, indicating the importance of controlling for family effects and population structure in Mendelian randomization studies.

## Results

We conducted a simulation to demonstrate how dynastic effects can affect Mendelian randomization estimates from unrelated individuals, and the methods that can be used to reduce these biases. Then we illustrate how these methods can be used with four empirical hypotheses examples using data from the HUNT and UK Biobank studies.

### Simulation Study

In simulations where the exposure does not have a causal effect on the outcome (**Figure 2**), the standard Mendelian randomization estimates were biased with high false discovery rates in the presence of dynastic effects (false discovery rate > 0.75 when the confounders were *C*_*x*_ and *C*_*y*_ = 0.1, *β*_*ux*_ *=* 0.1, n> 10,000). The sibling and trio methods were unbiased.

**Figure 2:**
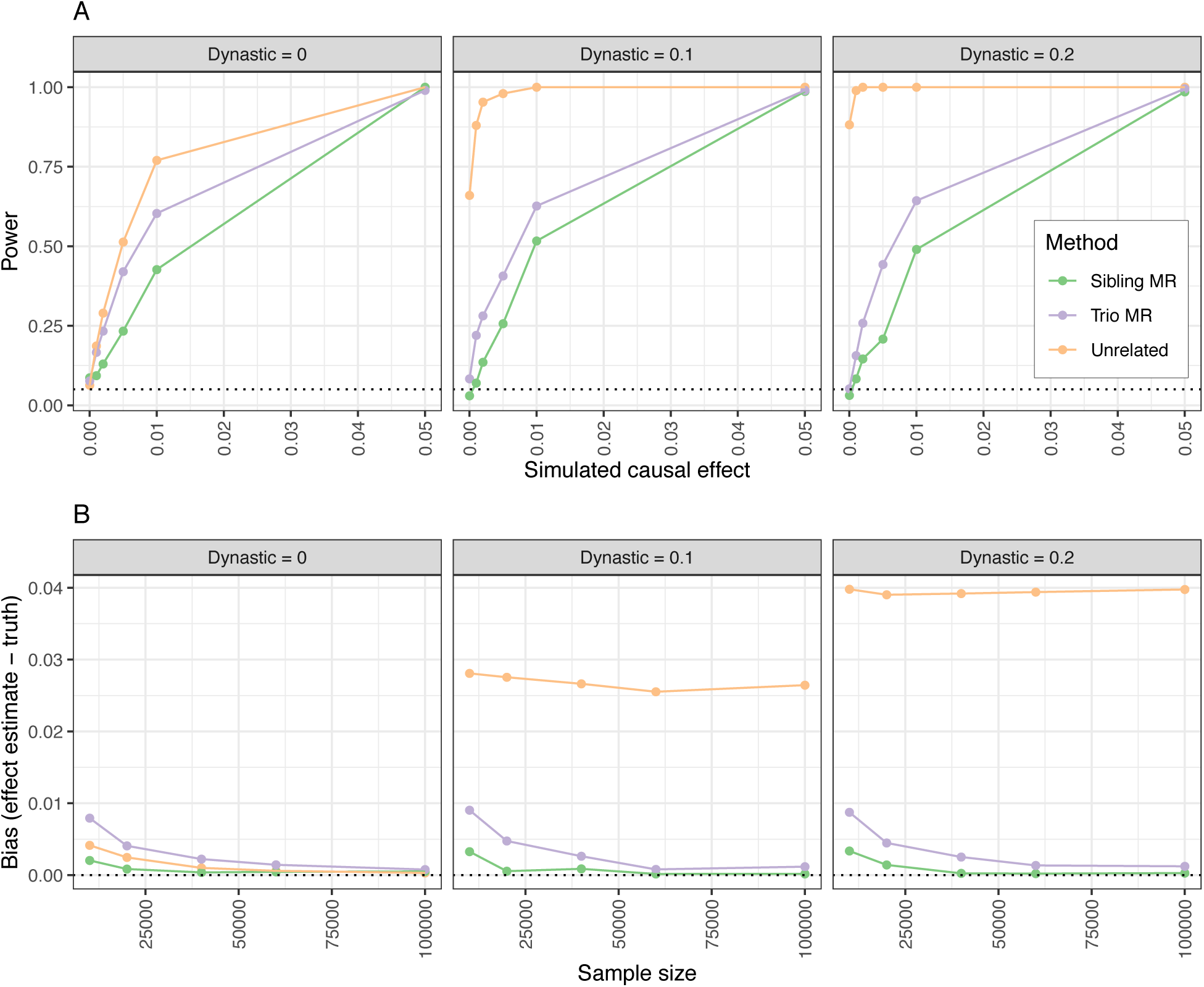
Estimated false discovery rate by power of the studies using different Mendelian randomization designs. A: SNP-exposure r^2^ = 0.05; sample size = 10000; simulation involves an influence of parental exposure influencing child’s confounder, which explains 10% of variance in child exposures and outcomes. For a simulated causal effect = 0, we expect the false discovery rate to be 0.05. B: Estimated bias by sample size using different Mendelian Randomization designs. The simulations are similar to panel (A) but allow sample size to vary and fixing the causal effect of an exposure x on an outcome y to 1% of variance explained. The bias in within-family Mendelian randomization estimates is small unless the dynastic effects are very small, or the number of observations modest.

Where we simulated the exposure to have a causal effect on the outcome Mendelian randomization using unrelated individuals had the highest power (**Figure 2**). However, the sibling and trio design also performed well with larger sample and effect size (power > 0.9 when sample sizes >= 10,000, dynastic effect <= 0.2, effect size = 0.05). The within-family models were substantially less powerful than standard Mendelian randomization using unrelated individuals; as usual, controlling bias comes at a cost.

### Empirical study

Participants with higher BMI were more likely to have diabetes: each 1kg/m^2^ increase in BMI was associated with a 0.60 (95%CI: 0.55 to 0.65, p-value<1.2×10^-136^) percentage point increase in the diabetes risk. These differences were modestly attenuated after including a family fixed effect (0.46, 95%CI: 0.40 to 0.52, p-value=8.5×10^-52^). The Mendelian randomization estimate using unrelated individuals suggested that each unit increase in BMI increased the risk of having diabetes by 0.82 (95%CI: 0.71 to 0.93, p-value=3.3×10^-50^) percentage points. This estimate remained after allowing for the fixed effects of family (1.01 percentage point increase per 1kg/m^2^ increase in BMI, 95%CI: 0.58 to 1.44, p-value=3.3×10^-06^). The summary data Mendelian randomization analysis allowing for family effects estimates were similar (0.75 percentage point increase per 1kg/m^2^ increase in BMI, 95%CI: 0.38 to 1.13, p-value=7.6×10^-05^, p_diff unrelated_= 0.74). On average the associations of the SNPs and BMI and diabetes were similar before and after allowing for a family fixed effect, falling 7% (95%CI: −5% to 20%, p-value=0.26) and increasing 11% (95%CI: −17% to 40%, p-value=0.42) respectively.

Participants with higher BMI were more likely to have high blood pressure; each 1kg/m^2^ increase in BMI was associated with a 2.63 (95%CI: 2.54 to 2.72, p-value<1×10^-300^) percentage point increase in high blood pressure risk. This association did not attenuate after including a family fixed effect (2.42, 95%CI: 2.30 to 2.54, p-value<1×10^-300^). The Mendelian randomization estimate using the sample of unrelated individuals suggested that each unit increase in BMI increased the risk of having high blood pressure by 1.59 (95%CI: 1.34 to 1.83, p-value=1.3×10^-36^) percentage points. The Mendelian randomization estimate was similar after allowing for a family fixed effect (1.13 percentage point increase per 1kg/m^2^ increase in BMI, 95%CI: 0.04 to 2.21, p-value=0.04). The summary data Mendelian randomization estimates were similar (0.76 percentage point increase per 1kg/m^2^ increase in BMI, 95%CI: −0.19 to 1.70, p-value=0.12, p_diff unrelated_= 0.10). On average the associations of the SNPs and high blood pressure fell by 51% (95%CI: 23% to 80%, p-value=0.0006) after allowing for family fixed effects.

Taller participants were more educated; each 10cm increase in height associated with an additional 0.45 (95%CI: 0.43 to 0.48, p-value=9.0×10^-307^) years of education (**Figure 3**). This association was attenuated after including a family fixed effect (0.22, 95%CI: 0.18 to 0.26, p-value=4.6×10^-25^). The Mendelian randomization estimate using the sample of unrelated individuals implied that each 10cm increase in height caused an increase of 0.17 (95%CI: 0.14 to 0.20, p-value=8.5×10^-26^) years of education. After allowing for a family fixed effect, the Mendelian randomization estimate was greatly attenuated suggesting little causal effect of height on education (mean difference per 10cm increase in height: 0.002, 95%CI: −0.13 to 0.13, p-value=0.98). When we used two sample Mendelian randomization by estimating the SNP-exposure and SNP-outcome associations in different samples (split sample)^35,36^ and then meta-analysed, there was little evidence of a causal effect of height on education (mean difference per 10cm increase in height=0.009, 95%CI: −0.11 to 0.13, p-value=0.87, p_diff unrelated_= 0.008). On average, the associations of these SNPs and height and education fell by 18% (95%CI: 14% to 22%, p-value=8.5×10^-24^) and 61% (95%CI: 49% to 73%, p-value=1.5×10^-21^) after allowing for family fixed effects respectively.

**Figure 3:**
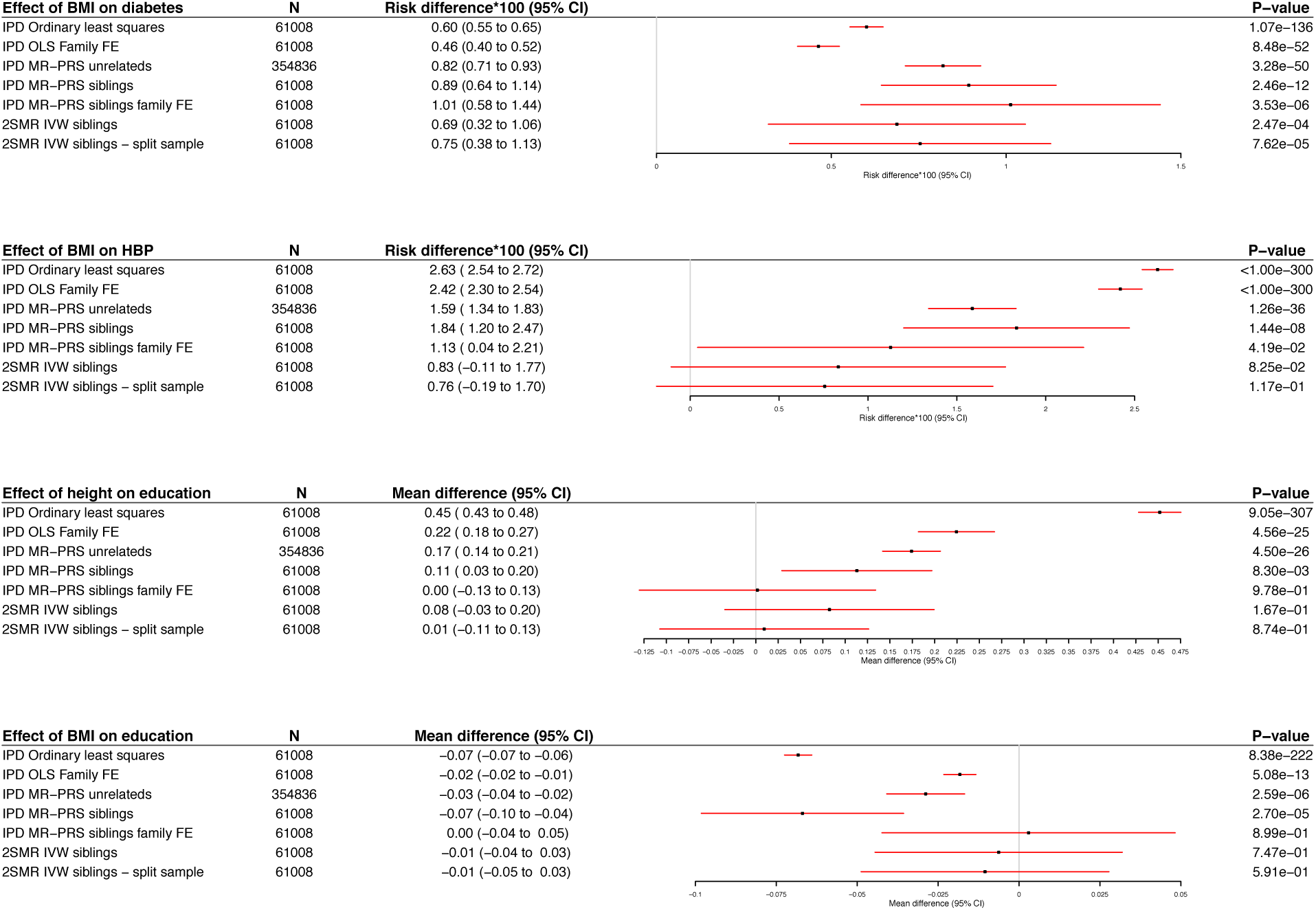
Estimates of the effect of BMI on self-reported diabetes and high blood pressure and height and BMI on educational attainment using ordinary least squares, Mendelian randomization in unrelated individuals and samples of siblings. All methods were consistent with BMI increasing diabetes and high blood pressure risk. Being taller and slimmer were phenotypically associated with higher educational attainment. However, the effects of height and BMI on educational attainment were attenuated but still detected by Mendelian randomization estimates using unrelated individuals from UK Biobank and HUNT. The effects were eliminated after allowing for a family effect using individual-level or summary data Mendelian randomization.

On average, participants with higher BMI were less educated: each 1kg/m^2^ increase in BMI was associated with 0.07 fewer years of education (95%CI: 0.06 to 0.07, p-value=8.4×10^-222^, see **Figure 3**). This association was attenuated after including a family fixed effect (0.02, 95%CI: 0.01 to 0.02, p-value=5.1×10^-13^). The Mendelian randomization estimate using the sample of unrelated people implied that each unit increase in BMI decreased years of schooling by 0.03 (95%CI: 0.02 to 0.04, p-value=2.6×10^-06^). This effect was eliminated after allowing for a family fixed effect, suggesting little causal effect of BMI on educational attainment (mean difference per 1kg/m^2^ increase in BMI= 0.00, 95%CI: −0.04 to 0.05, p-value=0.89). Again, the effect was also largely attenuated when we used two sample summary data approaches. Using separate samples to estimate the SNP-exposure and the SNP-outcome associations allowing for family fixed effects, there was little evidence of an effect of BMI on educational attainment (mean difference per 1kg/m^2^ increase in BMI=-0.01, 95%CI: −0.05 to 0.03, p-value=0.59, p_diff unrelated_= 1.6×10^-03^). On average, the association of the 69 BMI SNPs and education fell by 65% (95%CI: 34% to 76%, p-value=1.8×10^-06^) after allowing for family fixed effects. These results suggest that the estimates in unrelated individuals may be due to dynastic effects or assortative mating. We found little evidence of heterogeneity between the two sample Mendelian randomization estimates from UK biobank and HUNT, except for the effect of BMI on diabetes (p-value=0.027).

We investigated whether our results could be explained by pleiotropy using the weighted median, weighted modal and MR-Egger estimators. These summary data Mendelian randomization estimators use estimates of the SNP-exposure and SNP-outcome associations to estimate the effect of the exposure on the outcome. These estimators are robust to a number of forms of pleiotropy. There was little evidence of differences between the IVW and pleiotropy robust methods, pleiotropy from the MR-Egger intercept, or heterogeneity across the studies (**Supplementary Figure 3**).

## Discussion

We have presented within-family methods for Mendelian randomization and demonstrated how confounding due to family structure can bias Mendelian randomization studies using unrelated individuals. The simulations illustrated how bias occurs even if the phenotype of interest has no direct causal effect on the outcome, and these effects can theoretically cause false positive findings. The simulations further demonstrated how family structure can be exploited to control for these effects either using samples of siblings or parent-offspring trios. However, estimates from within-family Mendelian randomization are less precise than estimates using unrelated individuals, which is consistent with those seen for allelic association.^37–39^. Furthermore, in practice, there are fewer relatives than unrelated people in most studies. In samples from HUNT and UK Biobank, we investigated the impact of family structure on four empirical examples; the effects of BMI on the risk of diabetes and high blood pressure and the effects of height and BMI on educational attainment. We found that the effects of BMI on the risk of diabetes and high blood pressure were less precise, but similar when allowing for family effects. Conversely, the effects of height and BMI on educational attainment were almost entirely attenuated after allowing for family fixed effects.

A substantial literature has used Mendelian randomization and samples of unrelated individuals to establish that BMI increases the risk of diabetes and hypertension later in life.^40^ Our results suggest that confounding due to family structure is unlikely to be generating these results, and that they are more likely to be due to a causal effect of BMI on individual’s risk. Behavioural geneticists have used longitudinal data from samples of twins to understand how different family members affect each other over time.^41,42^ Other studies have used animal models to investigate how “social genetic effects” (i.e. indirect or dynastic effects) can affect health outcomes.^43^ A rich literature has established that height and BMI are respectively positively and negatively associated with educational attainment and socioeconomic position.^44–46^ Consistent with our results, previous studies using twin data have indicated that the relationship between height and educational attainment is likely to be due to non-genetic shared familial factors.^47,48^ These findings raise questions about whether height and BMI have causal effects on socioeconomic outcomes later in life.^49–51^ However, larger studies of related individuals with information on socioeconomic outcomes, such as income and occupation later in life are required to provide definitive evidence about the consequences of height and BMI later in life.

In general, within-family Mendelian randomization estimates are less precise than estimates from samples of unrelated individuals. Thus, within-family estimates of a specific association can be considered more robust, but less efficient estimates. Therefore, if there is evidence of differences between the estimates, then generally, the more imprecise but less biased within-family estimates should be preferred. Our estimates of the effect of height and BMI on educational attainment are an example of this situation. If there is little evidence of differences between estimates using unrelated and those allowing for family effects, then the former estimates should be preferred. Our estimates of the effect of BMI on risk of diabetes and high blood pressure are an example of this situation. This is analogous to comparing instrumental variable estimates to multivariable adjusted estimates.^3^ While allowing for family fixed effects or using difference estimators will account for dynastic effects or assortative mating, these methods will not address bias due to violations of the third Mendelian randomization assumption (exclusion restriction). This assumption is that the SNPs have no direct effect of the SNPs on the outcome (i.e. no pleiotropy). MR-Egger, weighted median and mode, or Lasso estimators are robust to various forms of violations of this assumption.^7–9,34^ It is trivial to use these estimators with the summary data methods we describe above and illustrate in **Supplementary Figure 3**. However, typically these estimators have lower power than the IVW estimator. The within-family summary data SNP-exposure and SNP-outcome associations, which allow for a family fixed effect, can be used with existing summary data estimators. Other proposed approaches allow for sophisticated control and estimation of pleiotropy and can trivially include family fixed effects,^52^ but again generally have lower power and require more data than Mendelian randomization approaches using allele scores or IVW. Therefore, given the statistical power of currently available samples of related individuals, investigators may be restricted to estimators that are robust to either family structure or pleiotropy, but not both. A further issue concerns residual population stratification in ancestrally heterogenous GWAS such as GIANT, which may bias SNP-phenotype associations for height, and affect analyses in other samples using SNPs identified in those GWAS.^22^ Within-family Mendelian randomization can control for residual population stratification.

Within-family estimates from samples of siblings, that allow for a family fixed effect are robust to biases due to dynastic effects, assortative mating and fine population structure.^1^ Of these approaches, the sibling design is potentially most useful because large amounts of such data are available through biobanks and family-based studies. Phenotypic similarity of siblings may reflect ‘passive’ sharing of environments or genes, or ‘active’ imitation or contrast effects arising from interaction between siblings.^53^ Contrast effects, which may inflate the estimated contribution of the nonshared environment in twin studies,^54^ can be mimicked by parental rating bias.^55,56^ However, for biological phenotypes where rating bias is not a concern, Mendelian randomization could be used to study the influence via imitation or contrast of one sibling’s genotype on the other’s phenotype, sometimes called ‘social genetic effects’,^43^ thereby adding to work on dynamic interplay between siblings. ^41,42^

Dynastic effects and assortative mating may cause bias in GWAS.^14^ If a GWAS is aiming to estimate the causal effect of variants on a given phenotype, then samples of unrelated individuals may produce biased estimates and potentially spurious findings. Future studies could re-run GWAS on a full range of traits on samples of siblings allowing for family fixed effects. This approach would also address concerns about residual population stratification in GWAS, which may bias SNP-phenotype associations in GWAS especially of heterogeneous ancestry.^22^ However, to detect genetic variants that explain 0.1% of the variance of either the offspring or maternal effects (i.e. a 2 df test) will require sample sizes of 50,000 mother-offspring pairs to detect genome-wide significant associations (α = 5×10^-08^). Sample sizes of around 10,000 will be required to partition known loci of similar size to the above into maternal and/or offspring genetic effects (α = 0.05).^57^ This sample size would provide valuable information about which phenotypes are likely to be most strongly affected by dynastic effects and assortative mating. It is likely that many, particularly biological, traits are relatively unaffected by these effects and thus GWAS results for these traits are unlikely to be biased due to these factors. Recent GWAS of social traits such as education reported the attenuation after allowing for family effects in their estimates in small samples.^30^ Further work in this area should include estimating the consequences of family structure for GWAS and Mendelian randomization estimates.

## Conclusions

Family structure can cause bias in Mendelian randomization studies. We found differences between estimates from unrelated individuals and within-family estimates in simulations and empirical analysis. The causal estimates of the effect of height and BMI on educational attainment were almost entirely attenuated after allowing for family fixed effects. Within-family methods, either using individual-level, or summary data Mendelian randomization approaches can be used to obtain unbiased estimates of the causal effects of phenotypes in the presence of dynastic effects, assortative mating and population stratification.

## Methods

### Statistical models

We describe four methods of using family data for Mendelian randomization below. If there are only two siblings, the difference and family fixed effects methods are equivalent, see appendix for proof.

The model to be estimated can be described as:

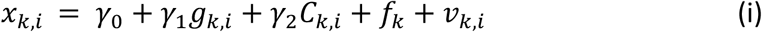

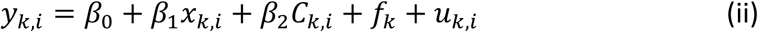

Where *y*_*k,i*_ and *x*_*k,i*_ are the outcome and exposure for individual *i* from family *k. g*_*k,i*_ is a set of genetic variants that are associated with the exposure. *f*_*k*_ is a family-level confounder, modelled in the empirical analysis via a family fixed effect (i.e. an indicator variable for each family). This accounts for all family-level confounders of the genetic variant-outcome association. Both *g*_*k,i*_ and *f*_*k*_ are functions of a family-level genetic component. *u*_*k,i*_ and *ν*_*k,i*_ are random error terms. *C*_*k,i*_ is a confounder of the association of the exposure and the outcome, *γ*_2_ and *β*_2_ indicate the effect of the confounder on the exposure and the outcome. *β*_1_ is the true causal effect of the exposure on the outcome which we wish to estimate. This model implies that Mendelian randomization using data from unrelated individuals would produce biased estimates of *β*_1_ due to the correlation between *g*_*k,i,j*_ and *f*_*k*_. The effect of the exposure on the outcome can be estimated using individual level data allowing for a family fixed effect, or summary level data using difference methods within families, or by allowing for a family fixed effect. We describe these approaches below.

#### (1) Siblings difference method

To apply Mendelian randomization to samples of siblings, effect estimates for the SNP-exposure association and SNP-outcome association are based on correlating the phenotypic divergence with the genotypic divergence within sibling pairs. Taking the difference between siblings removes the effect of the family-level confounder. For any pair of siblings within family *k*, indicated *k*, 1 and *k*, 2, the genotypic difference at genetic variant *j* is:

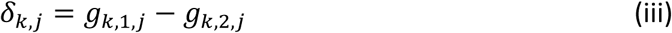

The association between the genotypic differences and phenotypic differences in the exposure, *x*, and outcome *y*, for SNP *j* can be estimated via:

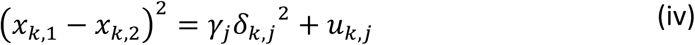

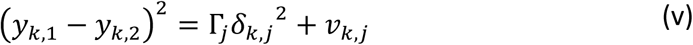

The estimated associations, *γ*_*j*_ and Γ_*j*_, can be used with any summary level Mendelian randomization estimator. Here we apply the inverse variance weighted (IVW) approach. Each pair of siblings can be included as a separate pseudo-independent pair.

#### (2) Family fixed effect with sibling data

Alternatively, we can estimate the associations using family fixed effects indicated by *f*_*k*_ for each family, which is equivalent to centring the data by subtracting the family mean.

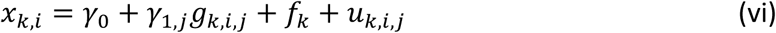

and

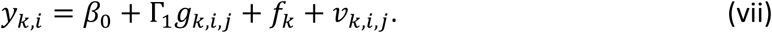

This estimator accounts for any differences between families, which includes any effect of assortative mating or dynastic effects common to all siblings by including a dummy variable for each family. This provides unbiased estimates of the SNP-exposure and SNP-outcome associations. These estimates can be used with standard summary data Mendelian randomization methods. The difference and family fixed effects methods are identical if there are only two siblings in each family. This fact follows from substituting equations iv and v into equations ii and iii and simplifying (see Appendix for proof). If there are more siblings, then the estimators are non-identical, but likely to be similar, see the appendix for further details. Cluster robust standard errors can be used to allow for clustering and relatedness within families.

#### (3) Adjusting for parental genotype with mother-father-offspring trio data

Finally, if data on mother-father-offspring trios are available, the estimates of the SNP-exposure and SNP-outcome associations for each child can be adjusted for their mother’s and father’s genotypes, indicated by *g*_*im,j*_ and *g*_*if,j*_ respectively^58^:

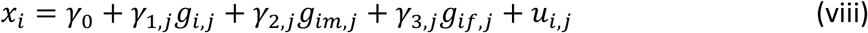

and

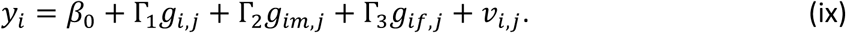

Again, these associations can be used to estimate the effect of the exposure on the outcome using summary data Mendelian randomization methods. It is possible to estimate the effect of foetal genotype on the exposure and outcome conditional on the mother and father genotype using summary data.^25,58^ The estimated causal effect can be biased if both the SNP-exposure and SNP-outcome associations are estimated in the same sample.^59^ This bias can be eliminated by splitting the sample and estimating the associations in separate samples.

#### (4) Two-stage least squares with sibling data

Many summary data methods assume no measurement error on the SNP-exposure association (NOME).^60^ This assumption may lead to underestimation of the standard error of the effect of the exposure on the outcome. Two-stage least squares can estimate the effect of the exposure on the outcome using the individual-level data from siblings. Estimators that use individual-level data can account integrate the estimation error from the SNP-exposure association. We used cluster robust standard errors that allow for clustering and relatedness within family. We used the commands xtivreg and plm.^61^

### Simulation of dynastic effects

We simulated a cohort consisting of pairs of unrelated mothers and fathers who had two offspring. All individuals had a genome of 90 SNPs. We set the distribution of identity by descent (IBD) across the 90 SNPs as *N*(0.5, 0.037) for each sibling pair, as per theory, because there are on average 90 recombination events separating human siblings. Hence, we assume that each SNP has an independent effect on the exposure.

We defined parents’ exposure and outcome by defining confounders *u*, exposure *x* and outcome *y*. A directed acyclic graph illustrating these relationships is shown in **Figure 1 (panel B)**. The confounder influences the parents’ exposure and outcome. The offspring have the same confounding structure, except the parent’s exposure affects their offspring’s outcomes, via a dynastic or ancestry effect. The genetic influence of each of the 90 SNPs on the exposure amounts to explaining *V*_*gx*_ of the variance in the exposure. We assumed no horizontal pleiotropy. All estimates assume *V*_*gx*_ = 0.1 and 90 independent causal variants (i.e. somewhat similar to GIANT results for BMI).^62^

To generate the phenotypes under a model of dynastic effects, the offspring outcome was influenced by both the offspring exposure and the parents’ exposures. In these simulations all phenotypes had mean of 0 and variance of 1. Differing strengths of dynastic effects, where the parental exposure influenced the offspring outcome were generated (*β*_*ux*_ = 0, 0.01, 0.02) under a set of models with a range of causal effects of the exposure on the outcome (*β*_*xy*_ = 0, 0.001, 0.002, 0.005, 0.01, 0.05). We calculated the false discovery rate (proportion of test with p-value < 0.05) for 100 iterations of each simulation using each of three methods: standard IVW as applied to one of each individual in a set of siblings (i.e. a sample of unrelated individuals), the within-family sibling design, and the within-family trio design. Finally, we calculated bias (estimated effect − simulated effect) for all three study approaches by simulated confounding (*C*_*x*_ and *C*_*y*_ = 0, 0.1, 0.2), dynastic bias (*β*_*Ux*_ = 0, 0.1, 0.2) and simulated causal effects (*β*_*xy*_ = 0, 0.001, 0.002, 0.005, 0.01, 0.05). The sample sizes were 10 000, 20 000, 40 000, 60 000 and 100 000 sibling pairs for all simulations.

### Empirical Analysis

To demonstrate the approach and assess potential dynastic effects, assortative mating, and residual population stratification, we conducted within-family Mendelian randomization using two illustrative examples in the Nord-Trøndelag Health Study (HUNT) and the UK Biobank.^63-65^ We estimated the effects of BMI on hypertension and diabetes and the effects of height and BMI on educational attainment. The effects of BMI on diabetes and hypertension have been well studied and provide a positive control.^40^ These effects on clinical outcomes experienced later in life are unlikely to be due to assortative mating or dynastic effects, because parents are less likely to assort on genetic liability for diabetes or high blood pressure. The genetic liability for these conditions was probably unknown when the couples were formed. Previous longitudinal and Mendelian randomization studies using unrelated individuals have suggested that height and BMI may affect educational attainment.^44,49^ Such an association, if causal, might be counteracted by changing educational policy. However, the association may be due to parents’ education, via dynastic effects or assortative mating, where more educated people select taller and slimmer partners. Assortative mating and dynastic effects can confound the association between genetic variants when data from the offspring generation are used. Therefore, the ratio of individual-level causal effects to family-level effects is likely to be higher for the effects of BMI on clinical end points than for the effects of height and BMI on education.

### HUNT descriptive data

There were 56,374 genotyped individuals in HUNT, including 11,448 families with at least two siblings comprising of 28,777 siblings (14,718 women) with complete data on genotype, height and education, diabetes and blood pressure. On average the participants in the full unrelated sample were 48.7 (SD = 15.1) years old, had a BMI of 26.3 kg/m^2^ (SD = 3.9), were 177.5cm tall (SD = 6.6) and 164.3cm tall (SD = 6.11) for men and women respectively, 2.5% of them had diabetes, and 42.5% had high blood pressure and had 12.0 years (SD = 2.1) of education. High blood pressure was defined as either currently taking anti-hypertensive medication or having systolic or diastolic blood pressure above 140mmHg or 90mmHg on average across up to three measurements in HUNT2. Descriptive statistics of the sample can be found in **Supplementary Tables 1** and **2**.

### UK Biobank descriptive data

There were up to 370,180 genotyped individuals in the UK Biobank, among whom were 16,847 families with at least two siblings, with 33,642 siblings (19,445 women) with complete data on genotype, height and education, diabetes and blood pressure. On average, the participants without siblings were 57.5 (SD = 7.4) years old, had a BMI of 27.4 kg/m^2^ (SD = 4.8), were 175.0cm tall (SD = 6.8) and 162.8cm tall (SD = 6.2) for men and women respectively, had 14.1 (SD = 2.3) years of education. 4.5% of them had diabetes and 54.0% had high blood pressure. High blood pressure was defined as either having a diagnosis of high blood pressure or having systolic or diastolic blood pressure above 140mmHg or 90mmHg respectively on average across up to two clinic measurements.

### Selection of genotypes for instruments

We selected 385 independent (r^2^<0.01 within 10,000kb) SNPs associated with height (p<5×10^-08^) from Wood et al. and 79 associated with BMI in Locke et al.^62,66^ Neither HUNT or UK Biobank were included in these studies. We clumped variants using the TwoSampleMR package.^67^ We harmonized the alleles’ effect sizes across the two samples and constructed weighted polygenic scores which were sums of the phenotype increasing alleles and weighted each variant by its effect on the phenotype in the published GWAS.

### The Nord-Trøndelag Health Study

The Nord-Trøndelag Health Study (HUNT) is a population-based cohort study. The study was carried out at four time points over approximately 30 years (HUNT1 [1984-1986], HUNT2 [1995-1997] and HUNT3 [2006-2008] and HUNT4 [2017-2019]). A detailed description of HUNT is available.^63^ We include 71,860 participants from HUNT2 and HUNT3 as they have been recently genotyped using one of three different Illumina HumanCoreExome arrays (HumanCoreExome12 v1.0, HumanCoreExome12 v1.1 and UM HUNT Biobank v1.0). For a flow chart of participants inclusion and exclusion from the study see **Supplementary Figure 1**. Imputation was performed on samples of recent European ancestry using Minimac3 (v2.0.1, http://genome.sph.umich.edu/wiki/Minimac3) from a merged reference panel constructed from the Haplotype Reference Consortium (HRC) panel (release version 1.1) and a local reference panel based on 2,202 whole-genome sequenced HUNT participants ^12-14^. Ancestry of all samples was inferred by projecting all genotyped samples into the space of the principal components of the Human Genome Diversity Project (HGDP) reference panel (938 unrelated individuals; downloaded from http://csg.sph.umich.edu/chaolong/LASER/)^15, 16^, using PLINK. We defined recent European ancestry as samples that fell into an ellipsoid spanning exclusively the European population within the HGDP panel. We restricted the analysis to individuals of recent European ancestry who passed quality control. Among these, 17,329 pairs of siblings comprising of 28,777 siblings, were inferred using KING^17^, where an estimated kinship coefficient between 0.177 and 0.355, the proportion of the genomes that share two alleles IBD > 0.08, and the proportion of the genome that share zero alleles IBD > 0.04 corresponded to a full sibling pair.

### Questionnaires, clinical measurements and hospitalizations

Participants attended a health survey which included comprehensive questionnaires, an interview and clinical examination. The participants’ height and weight were measured with the participant wearing light clothes without shoes to the nearest centimetre and half kilogram, respectively. Education was defined using the question “What is your highest level of education”. Participants answered one of five categories 1) primary school (=10 years), 2) high school for 1 or 2 years (=12 years), 3) complete high school (=13 years), 4) college or university less than 4 years (=16 years), and 5) college or university 4 years or more (=17 years). Participants with university degrees were assigned to 16 years of education, those who completed high school were assigned 13 years, those who attended high school for 1 or 2 years were assigned to 12 years, and those who only attended primary school were assigned to 10 years. Diabetes was defined using responses to the question “Have you had or do you have diabetes” which has high validity.^68^ High blood pressure was defined as those with systolic or diastolic blood pressure equal to or more than 140 or 90, respectively, or reported use of antihypertensive medication.

### Ethics

This study was approved by the Regional Committee for Medical and Health Research Ethics, Central Norway and all participants gave informed written consent (REK Central application number 2016/21296).

### The UK Biobank

The UK Biobank invited over 9 million people and sampled 503,317 participants from March 2006 to October 2010. The study sampled individuals from 21 study centres across Great Britain. A detailed description of the study can be read elsewhere.^64,65,69^ We included 33,642 related participants who were born in England to ensure that they experienced a similar school system. For a flow chart of participants inclusion and exclusion from the study see **Supplementary Figure 2**. The participants gave blood samples, from which DNA was extracted. Full details of the quality control process are available elsewhere. ^70^Briefly, we excluded participants who had mismatched genetic and reported sex, or those with non-XX or XY chromosomes, extreme heterozygosity or missingness. We limited the analysis to 11,554,957 SNPs on the Haplotype Reference Consortium (HRC) panel. We selected SNPs that were reported to be associated with height and BMI in GWAS.^62,66^ We used these GWAS because they did not include UK Biobank. We selected independent SNPs associated r^2^<0.01 within 10,000kb by selecting the SNP most strongly associated with the trait of interest. We created weighted allele scores for height and BMI by calculating a weighted average of the number of height or BMI increasing alleles each participant had.^59^

### Questionnaires, clinical measurements and hospitalizations

Weight (ID:21002) and standing height (ID:50) were measured using standardised instruments the baseline assessment centre visits. We defined education using the participants response to the touch screen questionnaires about their educational qualifications (ID= 6138). We defined educational attainment using the participants’ highest reported educational qualification at either measurement occasion. We assigned participants with university degrees to 17 years of education, those with professional qualifications such as teaching or nursing to 15 years, those with A-levels to 14 years, those with National Vocational Qualifications (NVQs), Higher National Diplomas (HNDs) to 13 years, General Certificate of Secondary Education (GCSEs), Certificate of Secondary Education (CSEs) or O-levels to 12 years, and those who reported no qualifications to 11 years, which was the legal minimum length of education for this cohort. Diabetes and high blood pressure were defined using responses to the self-reported touch screen questionnaire (ID=6150 and ID=2443). We used self-reported measures because measured blood pressure is affected by medication use. Missing values at the baseline visit were replaced by measures from subsequent visits if available.

### Covariates and standard errors

All analyses included age, sex, the first 20 principal components of genetic variation. Cluster robust standard errors were used to allow for heteroskedasticity and allow for clustering and relatedness across siblings within families. Inclusion of the covariates age, sex, and principal components did not meaningfully affect the within family estimates, as they are independent of genotype conditional on sibling genotype.

### Ethics

UK Biobank received ethical approval from the Research Ethics Committee (REC reference for UK Biobank is 11/NW/0382). This research was approved as part of application 8786.

### Empirical analyses

We compared seven empirical estimates of the BMI on self-reported diabetes and high blood pressure and the effects of the height and BMI on educational attainment:

1. **IPD Ordinary least squares**: The phenotypic association using ordinary least squares
2. **IPD OLS Family FE**: The phenotypic association using ordinary least squares allowing for a family fixed effect across siblings.
3. **IPD MR-PRS unrelateds**: An estimate of the effect using a sample of unrelated individuals using a polygenic risk score and two stage least squares.
4. **IPD MR-PRS siblings**: As with 2. above, but restricted to siblings.
5. **IPD MR-PRS siblings family fixed effects**: An estimate using individual level data from the full sample of siblings family fixed effects. The standard errors allow for clustering by family and integrate imprecision from the SNP-exposure association. The results from HUNT and UK Biobank were combined via meta-analysis. Estimated using the xtivreg and plm packages.^61^
6. **2SMR IVW siblings**: An estimate using SNP summary data for Mendelian randomization including family fixed effects. The SNP-exposure and SNP-outcome associations were estimated on the same sample. The SNP level estimates of the effect of the exposure on the outcome were estimated separately in HUNT and UK Biobank and the overall Mendelian randomization (Wald) estimates are calculated for each SNP. For each SNP the Wald estimate is the ratio of the SNP-outcome and SNP-exposure association. We combine the estimate using Inverse Variance Weighted (IVW) meta-analysis.
7. **2SMR IVW siblings - split sample**: As with 5. above, but the SNP-exposure and SNP-outcome associations were estimated in separate samples (i.e. split sample approach). The overall Wald estimates were combined via IVW meta-analysis as above.

### Sensitivity analyses

Finally, we tested for difference (p_diff unrelated_) between the Mendelian randomization estimates using the unrelated individuals and the summary data within-family estimates using the split sample approach (i.e. as in 6. above).^71^ We investigated whether our results could be explained by pleiotropy using the weighted median, weighted modal and MR-Egger estimators and the SNP-phenotype associations allowing for a family fixed effect.^8,10,72^ We used a split sample approach in which the SNP-exposure and SNP-outcome associations were estimated in separate samples. We estimated the percentage change in the SNP-phenotype coefficients with and without allowing for a family fixed effect.

## Data and code availability

The empirical dataset will be archived with the studies and will be made available to individuals who obtain the necessary permissions from the studies’ data access committees. The code used to clean and analyse the data and the SNP level summary statistics are available here: https://github.com/nmdavies/within_family_mr.

## Acknowledgements

Jonathan Beauchamp provided valuable comments and suggestions on an earlier draft of this manuscript. The Medical Research Council (MRC) and the University of Bristol support the MRC Integrative Epidemiology Unit [MC_UU_12013/1, MC_UU_12013/9, MC_UU_00011/1]. The Economics and Social Research Council (ESRC) support NMD via a Future Research Leaders grant [ES/N000757/1]. LDH is supported by a Career Development Award from the UK Medical Research Council (MR/M020894/1). This work is part of a project entitled ‘social and economic consequences of health: causal inference methods and longitudinal, intergenerational data’, which is part of the Health Foundation’s Efficiency Research Programme. The Health Foundation is an independent charity committed to bringing about better health and health care for people in the UK. The Nord-Trøndelag Health Study (The HUNT Study) is a collaboration between HUNT Research Center (Faculty of Medicine and Health Sciences, NTNU, Norwegian University of Science and Technology), Nord-Trøndelag County Council, Central Norway Regional Health Authority, and the Norwegian Institute of Public Health. The K.G. Jebsen Center for Genetic Epidemiology is funded by Stiftelsen Kristian Gerhard Jebsen; Faculty of Medicine and Health Sciences, NTNU; The Liaison Committee for education, research and innovation in Central Norway; and the Joint Research Committee between St. Olavs Hospital and the Faculty of Medicine and Health Sciences, NTNU. The genotyping in HUNT was financed by the National Institute of Health (NIH); University of Michigan; The Research Council of Norway; The Liaison Committee for education, research and innovation in Central Norway; and the Joint Research Committee between St. Olavs Hospital and the Faculty of Medicine and Health Sciences, NTNU. JK has been supported by the Academy of Finland (grants 308248, 312073). RMF and RNB are supported by Sir Henry Dale Fellowship (Wellcome Trust and Royal Society grant: WT104150). No funding body has influenced data collection, analysis or its interpretation. This publication is the work of the authors, who serve as the guarantors for the contents of this paper.

## Author contributions

GDS, NMD, KH and BMB obtained funding for this study. BMB, GH and NMD analysed and cleaned the data, interpreted results, wrote and revised the manuscript. GH ran the simulation study. ES provided the proof comparing the difference and fixed effects estimators, interpreted results and revised the manuscript. GDS conceived of the study, interpreted the results, wrote and revised the manuscript. All other authors interpreted the results and revised the manuscript.

## The Mendelian randomization within-family consortium

K.G. Jebsen Center for Genetic Epidemiology, Department of Public Health and Nursing, NTNU, Norwegian University of Science and Technology, Norway

Ben Brumpton, Gunnhild Åberge Vie, Johan Håkon Bjørngaard, Kristian Hveem, and Bjørn Olav Åsvold

Clinic of Thoracic and Occupational Medicine, St. Olavs Hospital, Trondheim University Hospital

Ben Brumpton

Medical Research Council Integrative Epidemiology Unit, University of Bristol, BS8 2BN, United Kingdom

Ben Brumpton, Eleanor Sanderson, Fernando Pires Hartwig, Sean Harrison, Yoonsu Cho, Laura D Howe, Amanda Hughes, Tim Morris, Laurence Howe, Oliver Davis, Claire Haworth, David Evans, George Davey Smith, Gibran Hemani, and Neil M Davies

Population Health Sciences, Bristol Medical School, University of Bristol, Barley House, Oakfield Grove, Bristol, BS8 2BN, United Kingdom

Eleanor Sanderson, Fernando Pires Hartwig, Sean Harrison, Yoonsu Cho, Laura D Howe, Amanda Hughes, Tim Morris, Laurence Howe, Oliver Davis, Claire Haworth, George Davey Smith, Gibran Hemani, and Neil M Davies

Postgraduate Program in Epidemiology, Federal University of Pelotas, Pelotas, Brazil

Fernando Pires Hartwig

Department of Biological Psychology, Vrije Universiteit Amsterdam, Amsterdam, The Netherlands

Dorret I Boomsma and Michel G Nivard

Netherlands Twin Register, Vrije Universiteit, Amsterdam, the Netherlands.

Dorret Boomsma

Department of Complex Trait Genetics, Center for Neurogenomics and Cognitive Research, Amsterdam Neuroscience, Vrije Universiteit, Amsterdam, the Netherlands

Philipp D Koellinger

Department of Economics, VU University Amsterdam, Amsterdam, the Netherlands

Philipp D Koellinger

German Institute for Economic Research, DIW Berlin, Berlin, Germany

Philipp D Koellinger

Nic Waals Institute, Lovisenberg Diaconal Hospital Spångbergveien 25, 0853 Oslo, Norway

Alexandra Havdahl

Department of Mental Disorders, Norwegian Institute of Public Health, Sandakerveien 24C, 0473 Oslo, Norway

Alexandra Havdahl

The University of Melbourne, 207 Bouverie St, Carlton, Victoria, 3010, Australia

John Hopper

Virginia Institute for Psychiatric and Behavior Genetics, Virginia Commonwealth University, Richmond, Virginia

Michael Neale

Karolinska Institutet, Dept of Medical Epidemiology and Biostatistics, Stockholm, Sweden

Nancy L Pedersen and Sara Hägg

Department of Psychology, University of California Riverside, Riverside, CA, USA Chandra Renyolds

Department of Psychology and Population Research Center, University of Texas at Austin

Elliot M Tucker-Drob, Andrew Grotzinger and K Paige Harden

Center for public health genomics, Department of public health sciences, University of Virginia, Charlottesville, VA, USA

Wei-Min Chen

Department of Biostatistics and Center for Statistical Genetics, University of Michigan, Ann Arbor, USA

Cristen Willer

Department of Internal Medicine, University of Michigan, Ann Arbor, MI, USA

Cristen Willer

Department of Human Genetics, University of Michigan, Ann Arbor, USA

Cristen Willer

University of Queensland Diamantina Institute, Translational Research Institute, Brisbane, Queensland, Australia

David M Evans

Department of Public Health, University of Helsinki, Helsinki, Finland

Jaakko Kaprio

Institute for Molecular Medicine Finland (FIMM), University of Helsinki, Helsinki, Finland

Jaakko Kaprio

Department of Endocrinology, St Olavs Hospital, Trondheim University Hospital, Trondheim, Norway

Bjørn Olav Åsvol

School of Psychological Science, University of Bristol, Bristol, United Kingdom

Claire Haworth

University of Edinburgh, Centre for Genomic and Experimental Medicine, Institute of Genetics and Molecular Medicine, University of Edinburgh, Crewe Road, Edinburgh, UK EH4 2XU

Archie Campbell

Institute for Molecular Medicine Finland (FIMM), P. O. Box 20 (Tukholmankatu 8) 00014 University of Helsinki

Teemu Palviainen

University of Minnesota, 75 East River Road, Minneapolis, MN 55455

William Iacono

The Danish Twin Registry, Department of Public Health, University of Southern Denmark, J.B. Winsloews Vej 9B, 5000 Odense C, Denmark

Kaare Christensen and Marianne Nygaard

Centre for Epidemiology and Biostatistics, Melbourne School of Population and Global Health, The University of Melbourne 207 Bouverie Street, Carlton, Victoria 3053, Australia

Shuai Li

Centre for Cancer Genetic Epidemiology, Department of Public Health and Primary Care, University of Cambridge, Strangeways Research Laboratory, Worts Causeway, Cambridge CB1 8RN, United Kingdom

Shuai Li

Italian Twin Register, Center for Behavioural Sciences and Mental Health, Istituto Superiore di Sanità Viale Regina Elena 299, I-00161 Rome, Italy

Maria Antonietta Stazi and Sonia Brescianini

Faculty of Psychology and IMIB-Arrixaca. University of Murcia. Campus de Espinardo. 30100 Murcia (Spain)

Juan R. Ordoñana

The University of Melbourne 207 Bouverie St, Carlton, Victoria, 3010, Australia

Maria Antontietta Stazi

GRIP, École de psychologie, Université Laval Pavillon Félix-Antoine-Savard, 2325 Allée des Bibliothèques, Québec, QC G1V 0A6, Canada

Michel Boivin

Karolinska Institutet Dept of Medical Epidemiology and Biostatistics, Stockholm, Sweden

Nancy L. Pedersen

University of Exeter, College of Medicine and Health, RILD Building, Barrack Road, Exeter, EX2 5DW

Rachel M Freathy and Robin Beaumont

QIMR Berghofer Medical Research Institute

Sarah Medland

University of Michigan, Ann Arbor, Michigan

Patricia Peyser

Department of Epidemiology, School of Public Health, University of Michigan, 1415 Washington Heights, Ann Arbor, MI 48109, USA

Lawrence Bielak

## Supplementary materials

### Equivalence of the first difference and fixed effects estimators when estimating within-family Mendelian randomization models

The general model as describe above is:

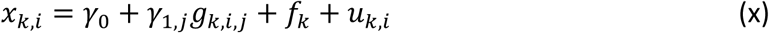

and

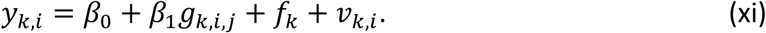

*f*_*k*_ is a family-level confounder, *U*_*I*_ is an individual-level confounder. *f*_*k*_ = *f*(*G*_*k*_, *NG*_*k*_) where *G*_*k*_ is a family-level genetic component and *NG*_*k*_ is a family-level non-genetic component, and *g*_*k,i,j*_ = *f*\*(*G*_*k*_). This setup means that Mendelian randomization using data from unrelated individuals is invalid due to the correlation between *g*_*k,i,j*_ and *f*_*k*_.

With two siblings the model can be written as:

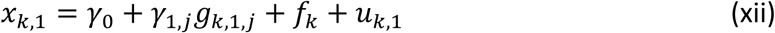

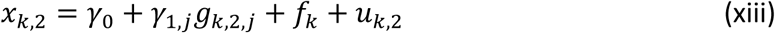

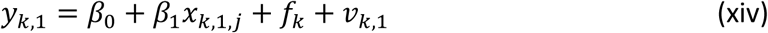

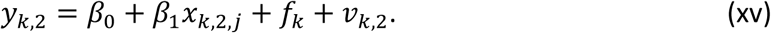

#### Fixed effects estimation

Fixed effects estimation estimates the model:

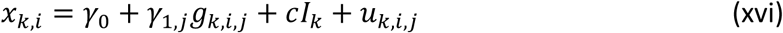

and

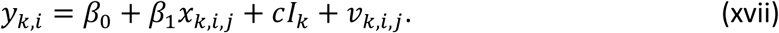

Where *I*_*F*_ is a set of family-level indicator variables that takes one value for each family (i.e. a set of extra indicator variables, the same size as the number of families in the model, each one of which takes 1 for one family and 0 for all other families). This model is estimated by taking the deviation from the family-level mean for each observation. Under this method of estimation our model becomes:

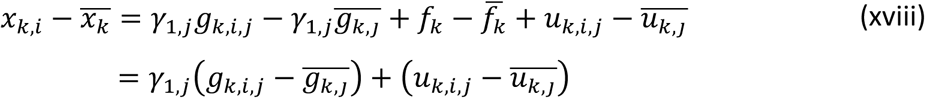

and

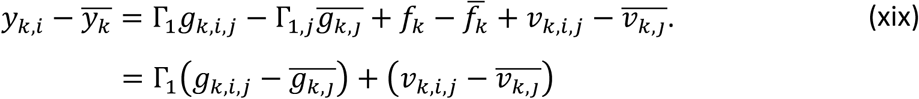

In a similar way to the First difference estimator, this estimator can now be consistently estimated using Mendelian randomization as 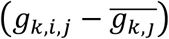 is independent of the family effect. This can be done by estimating:

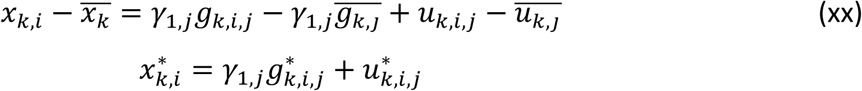

and

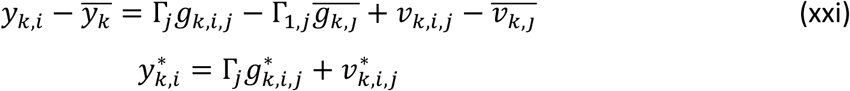

The two-sample MR estimator is then obtained from;

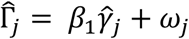

#### First difference (FD) estimation

A special case of the fixed effect estimator is the first difference estimator with two siblings. When the first difference is taken the model becomes:

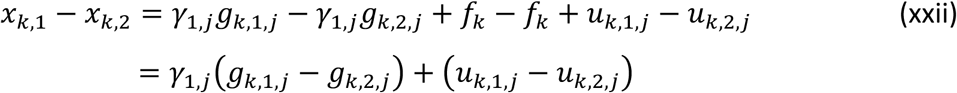

and

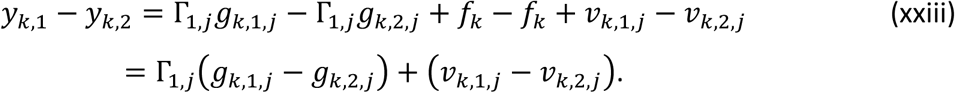

As (*g*_*k*,1_ − *g*_*k*,2_) is uncorrelated with (*u*_*k*,1,*j*_ − *u*_*k*,2,*j*_) this model can now be consistently estimated in a two-sample MR estimation by:

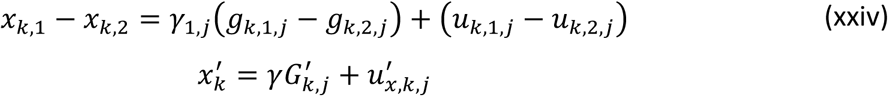

and

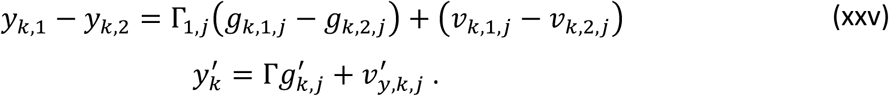

The two-sample MR estimation of 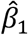 can then be calculated in the same way as above.

#### Equivalence of the first difference and fixed effects estimators

When there are exactly two individuals in each family these two methods of estimation will give the same result.

For *i* = 1 the variables in the fixed effects estimator is

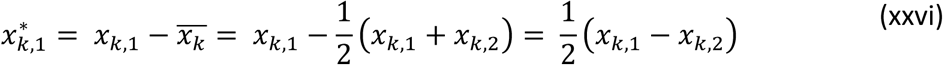

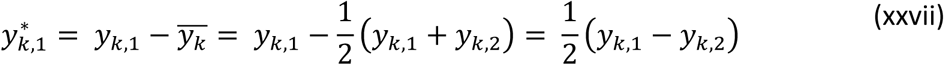

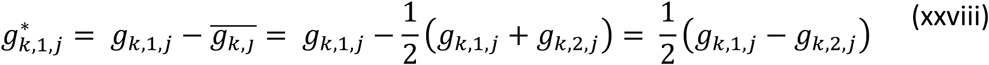

Therefore, the fixed effects estimator can be written as:

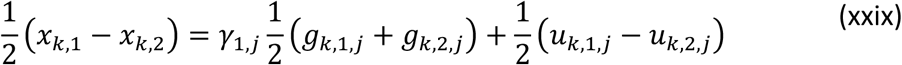

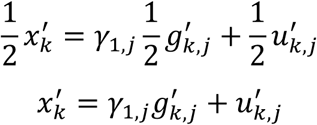

and

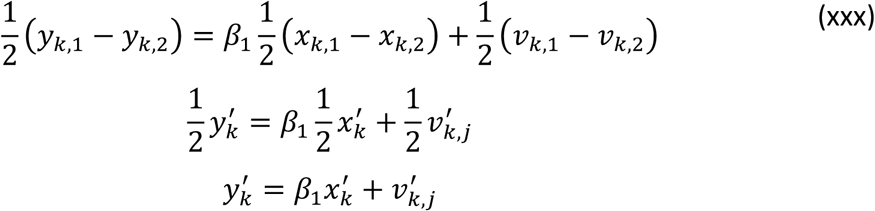

Therefore, the fixed effects estimator is equal to the first difference estimator with all of the variables included divided by 2. As this transformation is applied to every variable in the model it cancels out across the estimation and the estimated parameters will be the same as in the first difference model.

When there are more than two individuals in each family, the difference between these estimators depends on the error terms. If *v*_*k,i,j*_ is assumed to be uncorrelated between individuals in a family, then the fixed effects estimator is more efficient. However, if it is assumed to be dependent on the level taken by other family members (i.e.*v*_*k*,1,*j*_ = *v*_*k*,2,*j*_ + *ϵ*_*k*_ where *ϵ*_*k*_ is a randomly distributed variable) then the first difference estimator is more efficient.^73^

**Supplementary Figure 1:**
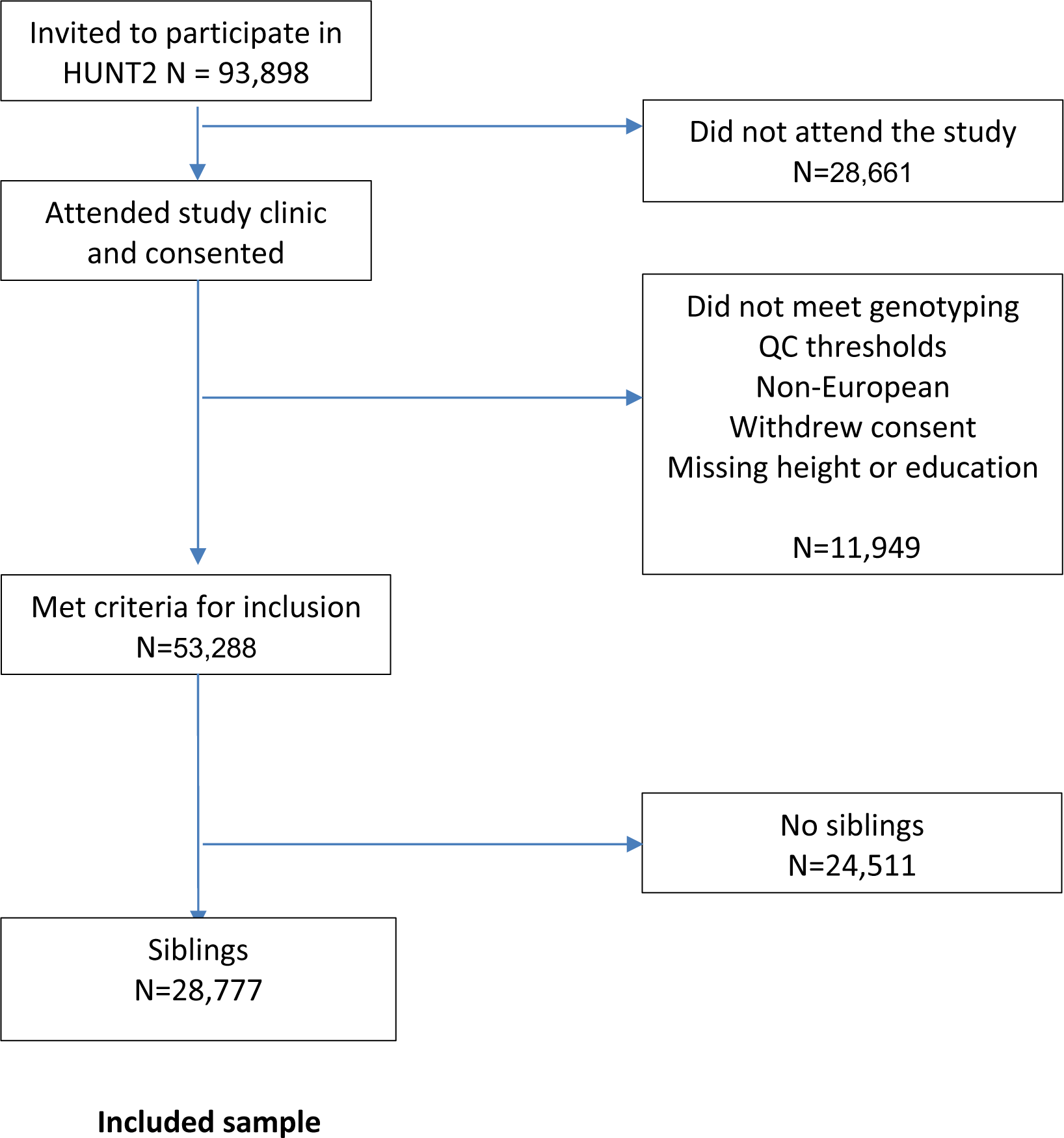
STROBE Flow chart representing inclusion into the study sample for HUNT2.

**Supplementary Figure 2:**
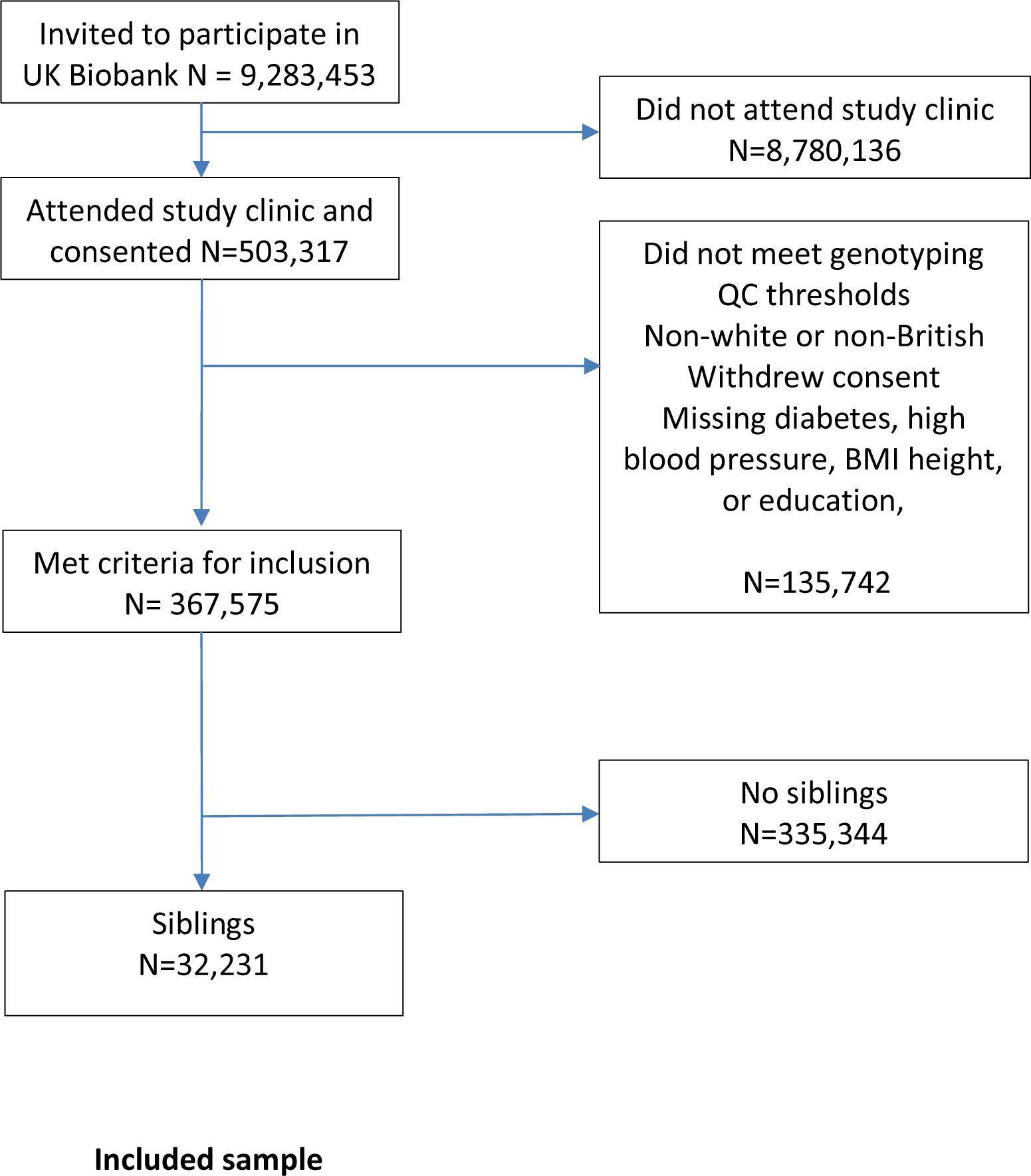
STROBE Flow chart representing inclusion and inclusion into the study sample for the UK Biobank.

**Supplementary Figure 3:**
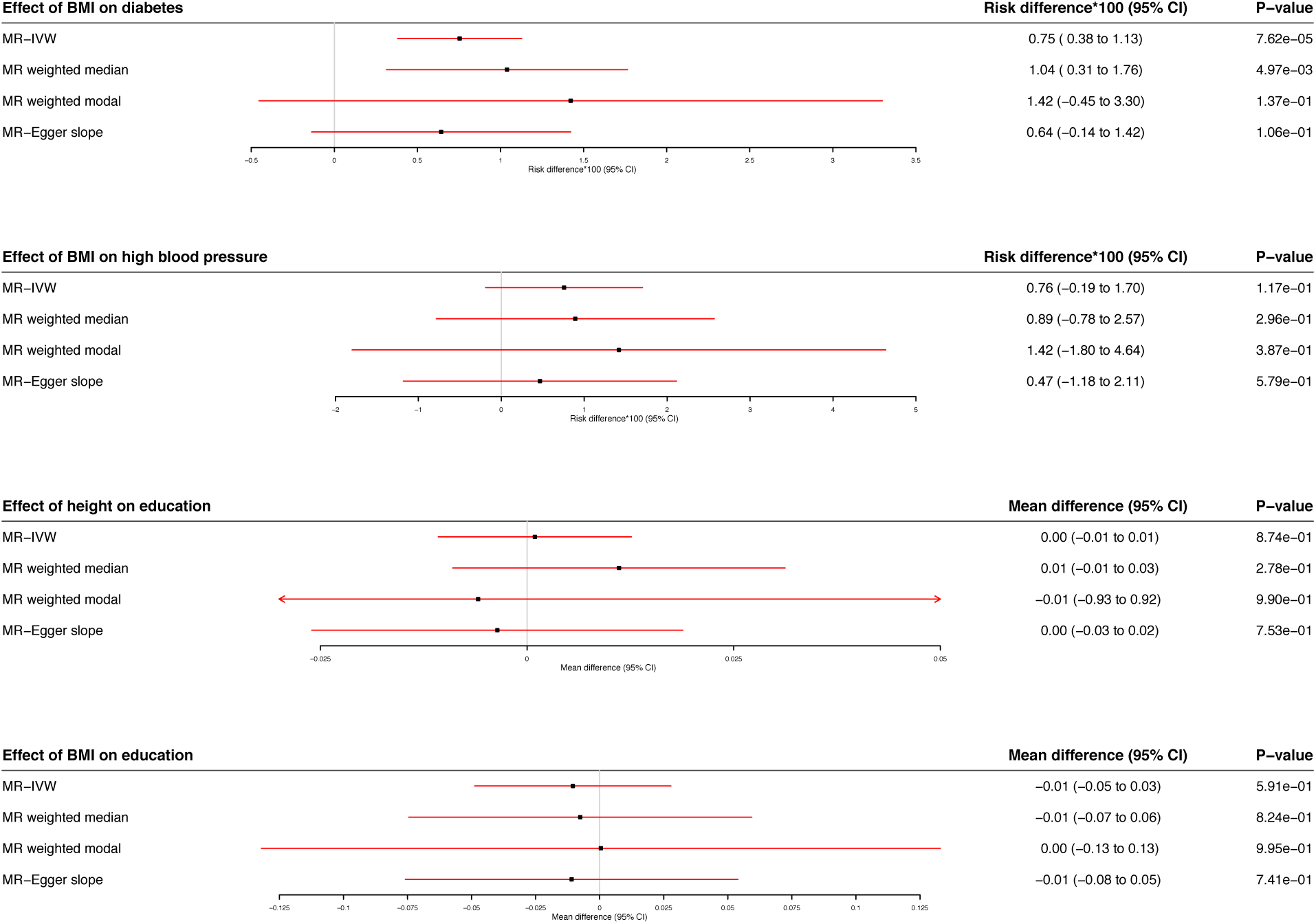
Estimates of the effect of BMI on self-reported diabetes and high blood pressure and height and BMI on educational attainment using Mendelian randomization in samples of siblings using inverse variance weighted (IVW), weighted median, weighted modal and MR-Egger. This analysis uses a split sample approach, in which the SNP-exposure and SNP-outcome associations are estimated in separate samples and allow for a family fixed effect. The weighted median, weighted modal and MR-Egger estimators are less precise than IVW. We found little evidence of pleiotropy using MR-Egger. There was little evidence of heterogeneity between HUNT and UKBB for any of the estimates. The MR-Egger intercepts for all outcomes found little evidence of directional pleiotropy (p>0.05), however this may be due to lack of power. The total analysed sample size in UK Biobank and HUNT was 61,008.

## Notes

https://github.com/nmdavies/within_family_mr

